# Implementation of a *Clostridium luticellarii* genome-scale model for upgrading syngas fermentations

**DOI:** 10.1101/2024.11.26.625427

**Authors:** William T. Scott, Siemen Rockx, Quinten Mariën, Alberte Regueira López, Pieter Candry, Ramon Ganigué, Jasper J. Koehorst, Peter J. Schaap

## Abstract

Syngas fermentation is a powerful platform for converting waste streams into sustainable carboxylic acid precursors for value-added biochemicals. Steel mills produce significant syngas, yet industrial microbial syngas valorization remains unrealized. The most promising syngas-converting biocatalysts consist of *Clostridia* species, such as *Clostridium kluyveri, Clostridium autoethanogenum*, and *Clostridium ljungdahlii. Clostridium luticellarii*, a recently discovered species, shares close phylogenetic ties with these organisms. Preliminary metabolic studies suggest its potential for syngas acetogenesis as well as chain elongation. In this study, we create *iSJ444*, a constraint-based metabolic model of *C. luticellarii* using *iHN637* of a close relative *C. ljungdahlii* as a starting point. Model predictions support hypothesized methanol and syngas pathways from the metabolic characterization studies; however, the use of propionate could not be accurately predicted. Thermodynamic Flux Analysis (TFA) reveals that *C. luticellarii* maintains stable energy dissipation across most reactions when exposed to varying pH, with significant increases observed in reactions associated with the Wood-Ljungdahl pathway (WLP), such as the HACD1 reaction, at higher pH (6.5), suggesting an adaptive role in energy management under neutral conditions. Flux sampling simulations exploring metabolic flux distributions show that *C. luticellarii* might fit into syngas fermenting platforms. In both cases, high hydrogen-to-carbon source ratios result in better production of (iso)butyrate and caproate. We present a minimal genome-scale metabolic model of *C. luticellarii* as a foundation for further exploration and optimization. Although our predictions of its metabolic behavior await experimental validation, they underscore the potential of *C. luticellarii* to enhance syngas fermentation platforms.

**Highlights:**

- *iSJ444* models *C. luticellarii* metabolism for syngas fermentation and chain elongation.
- Thermodynamic flux analysis (TFA) reveals adaptive energy balancing in pathways.
- Simulations highlight *C. luticellarii* as a producer of value-added biochemicals like butyrate, isobutyrate, and caproate
- Metabolic insights from *iSJ444* suggest efficient syngas conversion using varied substrates for industrial use.

## 1. Introduction

An important focus of modern biotechnological research is to understand and optimize microbial metabolic pathways for producing renewable and valuable chemicals, contributing to the advancement of a sustainable circular economy. These pathways enable the conversion of waste streams into useful products, often being present in a consortium of organisms to achieve enhanced product specificity or to produce compounds thermodynamically infeasible for a single species (Bekiaris and Klamt, 2021). Syngas fermentation stands out among various strategies due to its ability to convert industrial off-gases, such as those from steel mills, and gasified waste streams into a single, homogeneous feedstock that can be efficiently processed by a single organism to produce a diverse array of valuable products. Syngas, primarily composed of hydrogen, carbon monoxide, and carbon dioxide, can be utilized by anaerobic microorganisms, particularly acetogens, through the Wood-Ljungdahl pathway to produce chemicals like acetate, ethanol, and 2,3-butanediol (Gildemyn et al., 2017; Parera Olm and Sousa, 2021; Drake et al., 2008).

*Clostridium luticellarii* is a strictly anaerobic bacterium isolated from a mud cellar, notable for its butyrate production (Wang et al., 2015). The genome of *C. luticellarii* has been sequenced, revealing a close relationship to both *C. kluyveri* and *C. ljungdahlii* (Poehlein et al., 2018). A more comprehensive metabolic characterization by Petrognani et al. (2020) highlighted *C. luticellarii*’s potential to grow on methanol/acetate and H_2_/CO_2_, primarily producing butyrate, isobutyrate, and caproate. Additionally, they demonstrated its ability to synthesize valerate from propionate and methanol, a hypothesis previously proposed in an open-culture and autotrophically in a mono-culture (De Smit et al., 2019; Mariën et al., 2024). *C. luticellarii* thus possesses the dual capability of syngas fermentation into acetate (but not ethanol) and chain elongation pathways for both odd- and even-chained carboxylic acids. This rare combination of abilities is only observed in a few other organisms, such as *Eubacterium limosum* (Molitor et al., 2017), making *C. luticellarii* a promising candidate for inclusion in syngas upgrading systems.

Optimizing syngas fermentation for the production of valuable chemicals requires a deeper understanding of the complex anaerobic metabolic pathways involved. These pathways often exhibit flexibility, allowing metabolic shifts between products depending on environmental conditions such as substrate availability and pH. This metabolic adaptability is key to enhancing the production of higher-value chemicals like butyrate, caproate, and odd-chained carboxylic acids. While single strains, such as chain-elongating *Clostridium kluyveri*, can efficiently elongate carbon chains using ethanol and acetic or propionic acid (Baleeiro et al., 2019; Diender et al., 2019; Fernández-Naveira et al., 2019), incorporating such microbes into microbial consortia remains a potential strategy for leveraging their complementary metabolic capabilities. For instance, co-cultures of *C. kluyveri* with *Clostridium ljungdahlii* or *Acetobacterium wieringae* with *Anaerotignum neopropionicum* have successfully upgraded ethanol and acetate into more valuable products (Parera Olm and Sousa, 2023; Kim et al., 2019; Benito-Vaquerizo et al., 2020). However, the focus of future work lies in unraveling these metabolic shifts and understanding how environmental factors drive product specificity, which will inform the development of high-efficiency systems for industrial applications. Genetic engineering and synthetic co-cultures remain promising but secondary strategies for enhancing the metabolic versatility of syngas-fermenting systems.

GEnome-scale metabolic Modeling (GEM) is a vital tool in systems biology, offering a detailed framework for understanding the metabolic capabilities of organisms at the genome level. By reconstructing metabolic networks from genome annotations, GEM modeling enables the simulation of metabolic fluxes under various environmental and genetic conditions, which is crucial for elucidating complex metabolic pathways, predicting phenotypic outcomes of genetic modifications, and optimizing metabolic engineering strategies (Thiele and Palsson, 2010). Recent advances in GEMs have expanded their applications in diverse fields, including microbial biotechnology and bioprocessing (Atasoy et al., 2024).

GEMs have been instrumental in investigating metabolic processes in *Clostridia* species, particularly for the production of industrially relevant bioproducts such as short-chain fatty acids, organic acids, fuels and alcohols. For example, GEMs have been used to optimize propionate production in *Clostridium beijerinckii* by adjusting carbon fluxes (Diallo et al., 2019) and to enhance butyrate production in *Clostridium acetobutylicum* through pathway manipulations (Gallardo et al., 2016). Additionally, models have guided strain engineering strategies for improving the production of various target compounds (McAnulty et al., 2012; Scott Jr et al., 2020). Beyond organic acid production, GEMs have also been applied to explore biofuel pathways, such as in the genome-scale model of *Clostridium ljungdahlii* developed by Nagarajan et al. (2013) and the solventogenic metabolism model for *Clostridium beijerinckii* by Milne et al. (2011). These models have been pivotal in identifying metabolic bottlenecks and informing genetic modifications to enhance production efficiency across diverse bioprocesses.

In this study, we applied a constraint-based metabolic modeling approach to explore the potential of *Clostridium luticellarii* for syngas fermentation and direct chain elongation. We constructed the *iSJ444* model by adapting the BiGG (Biochemical, Genetic and Genomic) knowledge base model of *C. ljungdahlii* (iHN637) (Nagarajan et al., 2013), removing reactions without genetic evidence, and incorporating hypothesized pathways from De Smit et al. (2019) and Petrognani et al. (2020). The model was validated against the experimental results reported by Petrognani et al. (2020), and its behavior was further characterized by simulating growth and product spectrum under various syngas fermentation and methanol-chain elongation conditions. Thermodynamic flux analysis (TFA) Henry et al. (2007) was performed on the *C. luticellarii* GEM to investigate the role of pH in the production of butyrate, isobutyrate, and acetate. Additionally, TFA was used to examine energy dissipation across the network to identify potential metabolic bottlenecks impacting product formation efficiency.

## 2. Results

### 2.1. Building the *iSJ444* model: from template to functional GEM

The *iHN637* metabolic model of *C. ljungdahlii* was chosen as a template for building *iSJ444* due to its close phylogenetic relationship and the advantages of the standardized BiGG template. This approach facilitates model consistency, interoperability, and faster adaptation compared to *de novo* model construction. The BiGG format streamlines the process by providing a well-curated framework of reactions and metabolites, enhancing compatibility with existing databases and analysis tools. This reduces the need for extensive re-annotation or gap-filling, allowing for a focused effort on curating pathways specific to *C. luticellarii*. Consequently, using the BiGG template enabled the efficient and robust adaptation of the *iSJ444* model, leveraging the shared core metabolic framework while refining the organism-specific features.

The final *iSJ444* model consists of 444 genes, 735 reactions (including 100 exchange reactions), and 672 metabolites. Of these, 708 reactions (90% of all reactions in *iHN637*) were inherited, reflecting substantial overlap in core metabolism between the models. This overlap ensures that the well-established pathways from *iHN637* are retained, while the remaining 10% were specifically curated for *C. luticellarii* to incorporate unique metabolic traits and functionalities. Similar to *iHN637, iSJ444* includes only an extra- and intracellular compartment (no periplasm, which is present in some other metabolic models). This careful balance of adaptation and refinement underscores the utility of template-based modeling in achieving both accuracy and novelty for less-characterized organisms.

All reactions in the *iSJ444* GEM are mass balanced (except for the exchange reactions and the biomass function). While the original GEM sets all metabolite charges to 0, we have also created a version of *iSJ444* that incorporates metabolite charges using data extracted from the BiGG database. This updated model achieves 84.75% charge balance and 98.9% mass balance, as verified using MEMOTE. However, for consistency with the original modeling framework and to maintain alignment with the assumptions and datasets used in prior analyses, we proceeded with the version of *iSJ444* with metabolite charges set to zero for the main analyses presented in this study. The biomass function in the *iSJ444* GEM was adopted directly from *C. ljungdahlii* (*iHN637*) without modification, as defining a species-specific biomass function requires extensive experimental data. Although biomass composition can vary even among closely related species, using a function from a similar organism provides a reasonable approximation in the absence of precise data for *C. luticellarii*. In this study, the primary focus was on flux sampling to explore the range of possible metabolic flux distributions under different conditions, rather than optimizing for specific objectives such as biomass yield. Flux sampling allows a broader exploration of metabolic capabilities without the need for a predefined objective function. While the biomass reaction provides a theoretical framework for cellular growth, its role in this study was limited to serving as a reference for model validation rather than as an optimization target for flux predictions. A total of 532 of the 735 reactions (72%) have at least one gene assigned to them, slightly lower than the original *iHN637* model (78% coverage), partly because no genes are assigned to some of the added pathways. MEMOTE gives *iSJ444* a total score of 89%, with a consistency of 99.86%. This score is slightly higher than *iHN637* (98%), as some flux loops were removed. The annotation scores for metabolites, reactions, and genes are similar to those in *iHN637*. Changes were also made in the directionality of reactions (Table S2). For example, the reaction catalyzing the ligation of formate and tetrahydrofolate (THF) to formyl-THF (FTHFL*i*) is irreversible in *iHN637*, but is made reversible in *iSJ444*. This adjustment allows for ATP generation from methanol as a feedstock, as commonly reported for methylotrophic acetogens (Kremp et al., 2018; Kremp and Müller, 2021).

#### 2.1.1. Incorporation of chain elongation pathways in iSJ444

Two pathways were introduced in the *iSJ444* model (Figure 2), along with transport and exchange reactions for the newly introduced substrates and products of interest (methanol, isobutyrate, caproate, etc.). These pathways are based on the only two studies describing the metabolism of *C. luticellarii* (De Smit et al., 2019; Petrognani et al., 2020). None of the reactions added have genes assigned to them, as an analysis of the responsible genes was not carried out. However, the eggNOG orthology mapping may provide clues, as many *C. kluyveri* genes were identified as the closest orthologs. It is likely that *C. luticellarii* shares parts of its chain elongation genes with *C. kluyveri* (Table 1).

**Table 1.**
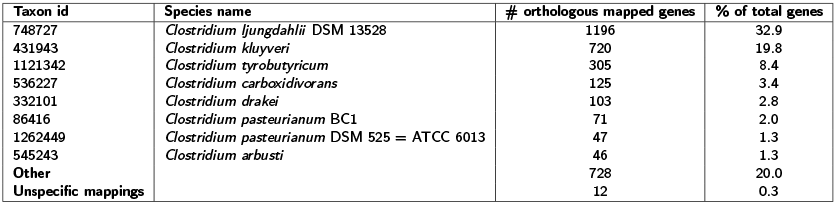
Species prevalence in eggNOG orthology mapping of *C. luti-cellarii*. Carried out on standard settings on the web service of eggNOG-mapper (Cantalapiedra et al., 2021). Unspecific mappings did not have a specific gene associated with them.

**Figure 1:**
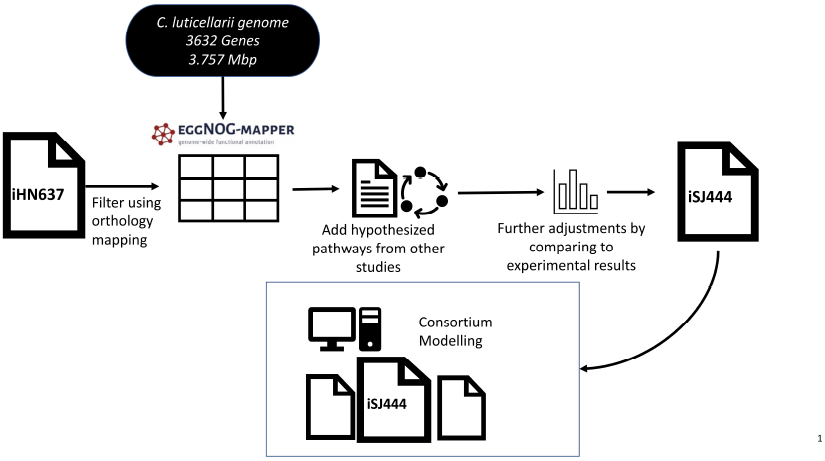
Setup of this study. The iHN637 model of *C. ljung-dahlii* was adapted to a *C. luticellarii* model (iSJ444) using an orthology gene mapping (eggNOG-mapper) and addition of hypothesized chain elongation-pathways from (De Smit et al., 2019; Petrognani et al., 2020). *iSJ444* can then be potentially joined with other preexisting metabolic models into consortia models and their potential for syngas recycling can be explored.

**Figure 2:**
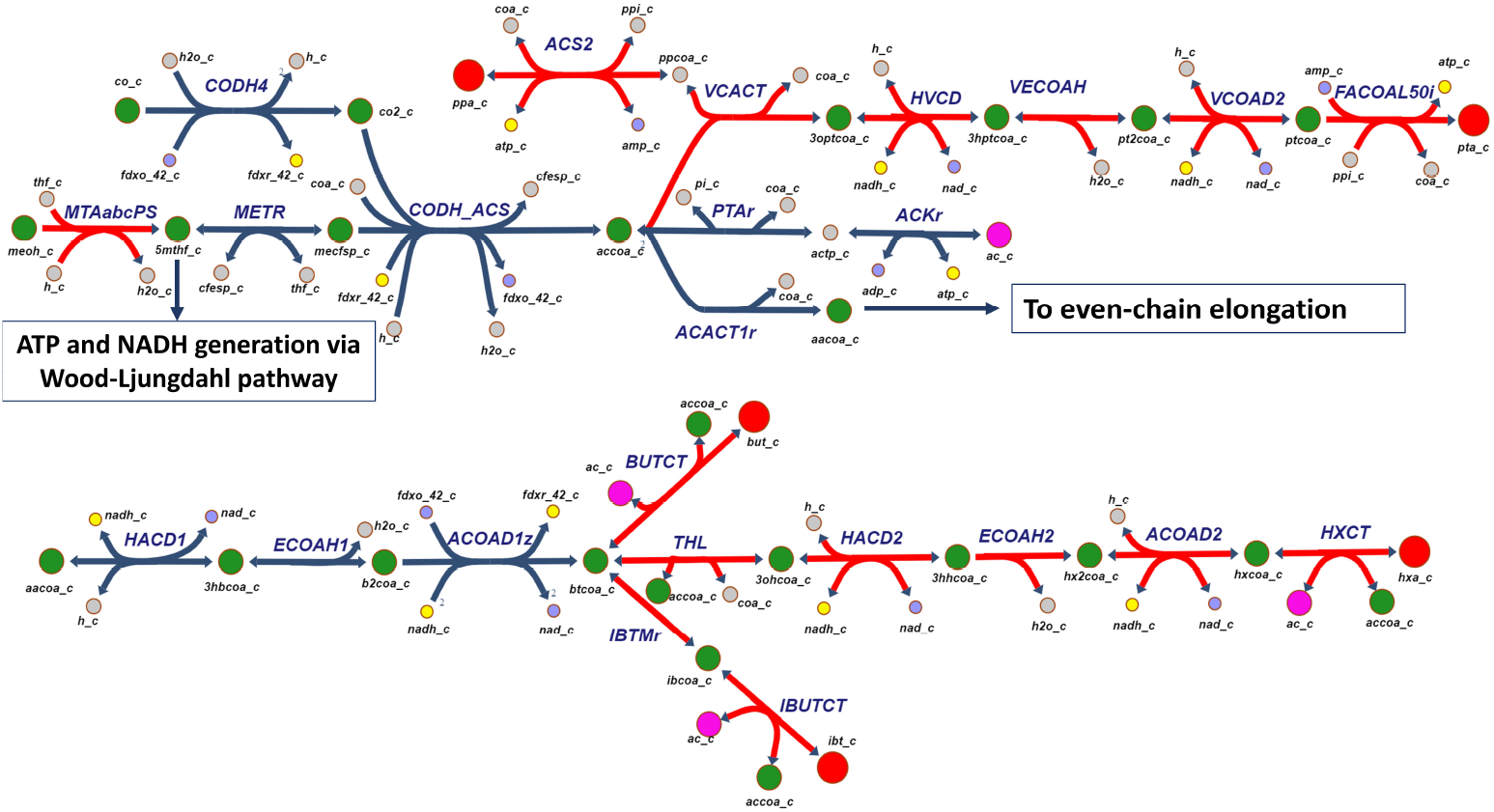
Hypothesized chain elongation pathways from (De Smit et al., 2019; Petrognani et al., 2020) were added to the *iSJ444* model. The starting steps of chain elongation were already present in *iHN637*, as well as the WLP, which, when growing on methanol, is used for ATP generation instead of carbon fixation. Reactions in blue were already present in *iHN637*. Red reactions are unique to *iSJ444*. Intermediate metabolites are green, energy-rich compounds (ATP, NADH, and reduced ferredoxin) are yellow, while their counterparts are light blue. The final products of interest are red, acetate is pink. Names of metabolites and reactions can be found in the abbreviations section. Figure is made using Escher (King et al., 2015).

Methanol is incorporated by the MTAabc system, a cobalamin-dependent system found in acetogenic bacteria (Kremp and Müller, 2021). The model simulates this through a pseudo-reaction that converts hydrogen, methanol, and tetrahydrofolate (THF) to 5-methyltetrahydrofolate (5mthf) and water. This intermediate is then funneled into the Wood-Ljungdahl Pathway (WLP), either towards CO_2_ to yield energy in the form of ATP and NADH, or directed towards acetogenesis and chain elongation.

### 2.2. Model validation: comparison to experimental results

The focus of this study was primarily on chain elongation and syngas metabolism, so these conditions were examined in greater detail. Predictions on various feedstocks were compared to experimental data to validate the model. Unless otherwise specified, experimental results referenced here are from Petrognani et al. (2020).

A comparison of substrates tested by Petrognani et al. (2020), on which *C. luticellarii* exhibited growth, is illustrated (see Figure 3). The modeled product spectrum on methanol and acetate closely matches the experimental results. The isobutyrate/butyrate/caproate production percentages are 41/40/11 and 43/48/8 for the model and experiment, respectively, when using flux sampling with minimal constraints (see Table S2) on metabolite uptake results in high standard errors. When butyrate is also supplied, both the model and experimental results show a shift in the product spectrum towards greater production of isobutyrate and caproate.

**Figure 3:**
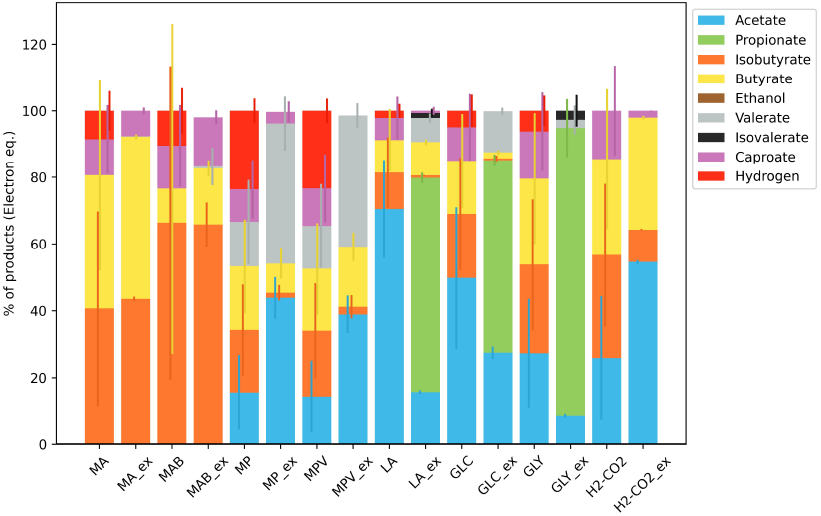
Product spectrum of the *iSJ444* model on different feedstocks, compared to experimental measurements by Petrognani et al. (2020). Bars show the average electron equivalent percentage (%) of the total spectrum of chain elongation products, as defined in the legend. Media names are explained in the abbreviations section. The suffix “_ex” indicates experimental results, while results without a suffix are model predictions. Error bars represent standard deviations. The simulation results were obtained using flux sampling with the Artificial Centered Hit-and-Run (ACHR) algorithm. The average electron equivalent was calculated by multiplying the flux of each product by its degree of reduction (see Table S1), followed by normalization against the total electron equivalents of all products.

The microbial growth on methanol with propionate as an electron acceptor showed greater divergence between model predictions and experimental results, both with and without valerate addition. The model predicted higher (iso)butyrate and caproate formation at the expense of valerate production. Similar over-predictions of these products were observed during growth on 80% H_2_ and 20% CO_2_, with modeled acetate production being half of the experimentally observed value: 26% versus 54%. Model predictions deviated most significantly from experimental measurements when grown on lactate, glucose, and glycerol. While propionate was experimentally observed to be the main fermentation product under these conditions, the model predicted no propionate formation and instead identified acetate as the major product (Figure S1). Additionally, (iso)valerate production was reported by Petrognani et al. (2020) as additional product, whereas the model predominantly suggested the formation of even-chained products.

### 2.3. Model performance across various substrates

#### 2.3.1. Growth on methanol and propionate

As earlier simulations showed a deviation from experimental results for growth on propionate, this was investigated further. When constraining the model to the methanol-propionate uptake ratio from the experiments (2:1) or (0.5 as presented in the figure), the model predicts a 1-to-1 propionate-valerate production ratio, as propionate can only be metabolized to valerate (Figure 4). This result contrasts with experimental findings, where the ratio is closer to 2-to-1 propionate uptake to valerate production. It is possible that under certain conditions, *C. luticellarii* oxidizes a fraction of the propionate to acetate, as observed in related cases of organic acid oxidation such as valerate to propionate or butyrate to acetate (Mariën et al., 2024), which could partially explain the higher acetate production observed.

**Figure 4:**
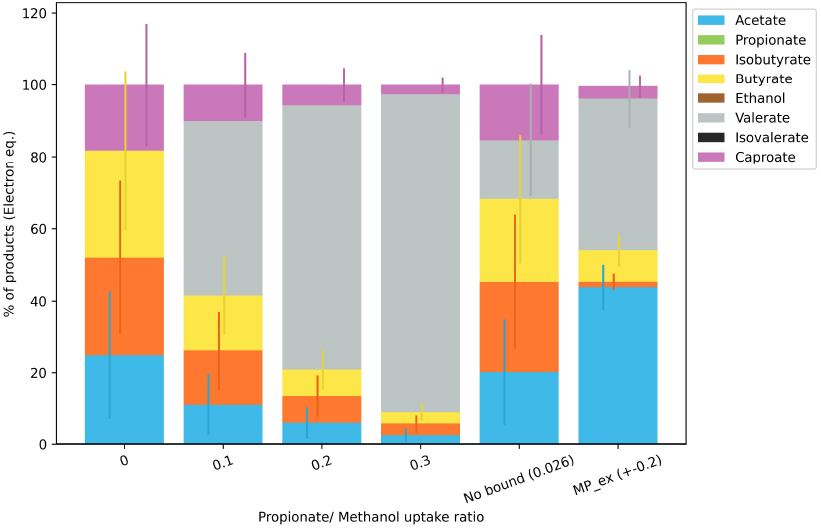
Modeled effect of uptake of propionate together with methanol. For the first 4 bars, propionate uptake was constrained to a specific ratio to methanol; in the ‘no bound’ condition, this was unconstrained. MP_ex refers to *in vitro* experimental data from Petrognani et al. (2020), where (+-0.2) indicates the ratio between propionate and methanol uptake as measured. As valerate is the only product propionate can be metabolized into, the electron equivalent of valerate production was approximately twice the equivalent of the consumed propionate (propionate has a degree of reduction of 14, valerate one of 26). H_2_ is also produced (see Figure S2), but is excluded from this graph as it was not measured in the *in vitro* experiments. Deviation bars indicate standard deviation.

#### 2.3.2. Influence of CO_2_ uptake on methanol product spectrum

Simulations on syngas and methanol showed no significant changes in the range or proportions of produced metabolites. In a potential industrial reactor design, *C. luticellarii* could be provided with syngas and syngas-derived methanol. Understanding what limits the production of chain elongation products is therefore valuable. Simulations indicate that increased CO_2_ uptake negatively affects the product spectrum; more acetate is produced at the expense of H_2_ and longer even-chain carboxylic acids (ECCAs) (Figure 5). CO_2_ uptake was not forced, so the uptake did not necessarily match the influx/availability, with the increase halting at around 7.0 mmol/gDW/h, along with the alteration in products. A significant reduction in H_2_ production is coupled with increased acetate as the final product. H_2_ production decreases from 28% of the electron equivalence of products to just 3% when comparing no CO_2_ uptake to the maximum unconstrained uptake. Conversely, acetate production increases from 17% to 63%. The mean total amount of chain elongation products decreases from 55% to 34%. This shift to the less desirable acetate production occurs just below a 2-to-1 methanol-CO_2_ uptake ratio.

**Figure 5:**
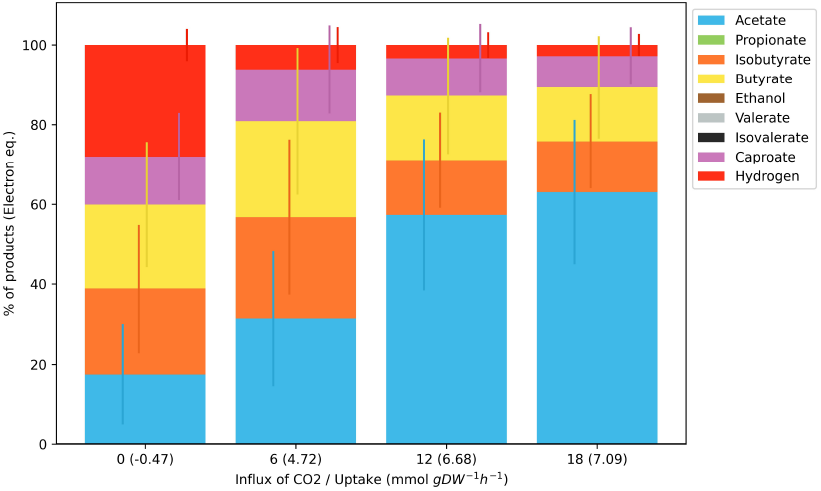
Simulated influence of CO_2_ uptake on *C. luticellarii* product spectrum during growth on methanol. The x-axis shows the maximum CO_2_ uptake flux (mmol/gDW/h) used in simulations, while numbers in parentheses represent the resulting median flux values derived from flux sampling. Methanol flux was fixed at 20 mmol/gDW/h. As more CO_2_ is consumed, H_2_ formation and chain elongation decrease, while acetate production increases. Deviation bars indicate the standard deviation of flux sampling results.

**Figure 6:**
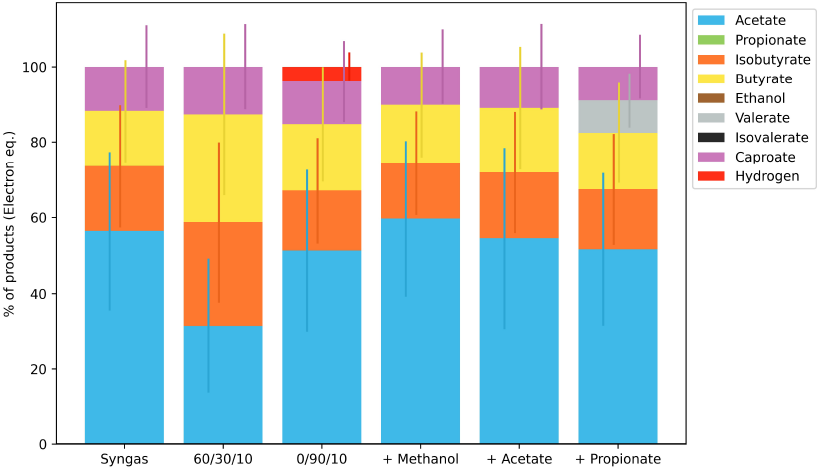
*iSJ444* model prediction of *C. luticellarii* product spectra on various syngas compositions and with the addition of trace feeds. x/x/x refers to H_2_/CO/CO_2_ ratios. + Methanol, acetate, and propionate were added at 5, 5, and 2 mmol/gDW/h influx of the respective metabolite in addition to syngas. Note the syngas composition here refers to 30/60/10 H_2_/CO/CO_2_ ratios, respectively. Deviation bars indicate standard deviation.

#### 2.3.3. Syngas fermentation and the addition of trace feeds

As the *iSJ444* model was designed to explore the syngas fermenting abilities of *C. luticellarii*, multiple simulations were conducted using different syngas compositions. The growth of *C. luticellarii* on syngas has not been extensively studied, so future experiments are needed to validate these predictions. In particular, growth on CO should be tested, as it is uncertain whether *C. luticellarii* can tolerate (high concentrations of) this toxic gas. CO can negatively affect bacterial growth even at micromolar levels (Mendes et al., 2021). While two CO-tolerant strains, *C. kluyveri* and *C. ljungdahlii*, are close relatives, recent studies indicate that *C. kluyveri* exhibits significant CO toxicity and only functions in a co-culture when CO partial pressures are kept low. Moreover, adaptive laboratory evolution (ALE) has been required to improve *C. kluyveri*’s tolerance to even moderate CO concentrations. Therefore, while *C. luticellarii* may share some tolerance mechanisms, it is premature to assume high CO tolerance without experimental validation (Diender et al., 2016; Mohammadi et al., 2012).

Results from simulations with various syngas compositions indicate no change in the product spectrum, except in cases with limited carbon availability in the form of CO or CO_2_, while maintaining constant energy availability through H_2_. In these cases, acetate production was approximately 31%, while in other media, it ranged from 51% to 60%. The model indicates the potential for *C. luticellarii* to grow using CO as the sole energy source. Simulations with additional acetate influx show no feasibility for extra acetate uptake by *C. luticellarii*, likely because acetate is already a primary end product of energy generation from syngas. Uptake of additional acetate would require further energy. The model showed unconstrained propionate uptake when fed syngas, which enabled valerate production. As indicated by previous simulations, the conversion of propionate in the current model is not energy-efficient, resulting in low uptake rates. Methanol addition did not affect the product spectrum on syngas.

### 2.4. Thermodynamic analysis of *C. luticellarii* metabolism under varying pH conditions

The Thermodynamic Flux Analysis (TFA) results, depicted in Figure 7, reveal how energy dissipation varies across the top 50 metabolic reactions of *Clostridium luticellarii* when exposed to pH levels of 5.5 and 6.5. For the majority of these reactions, dissipation changes were modest, typically remaining below 0.1 kJ/mol, suggesting that the organism’s metabolic network retains a considerable degree of stability, effectively maintaining its energy balance despite fluctuations in environmental pH. However, several reactions exhibited more pronounced shifts, indicating specific metabolic processes adjust their energy distribution in response to pH changes. These pH levels were selected to explore a hypothesis underscored in the experimental work of Mariën et al. (2024), which identified pH as a key factor influencing metabolic flux direction in *C. luticellarii*. Specifically, their findings showed that mildly acidic pH (≤ 5.5) stimulates the production of longer-chain products, such as butyric acid, while circumneutral pH (∼ 6.5) favors acetic acid production. By simulating energy dissipation at these pH levels, we aimed to provide a thermodynamic perspective on the metabolic shifts observed experimentally.

**Figure 7:**
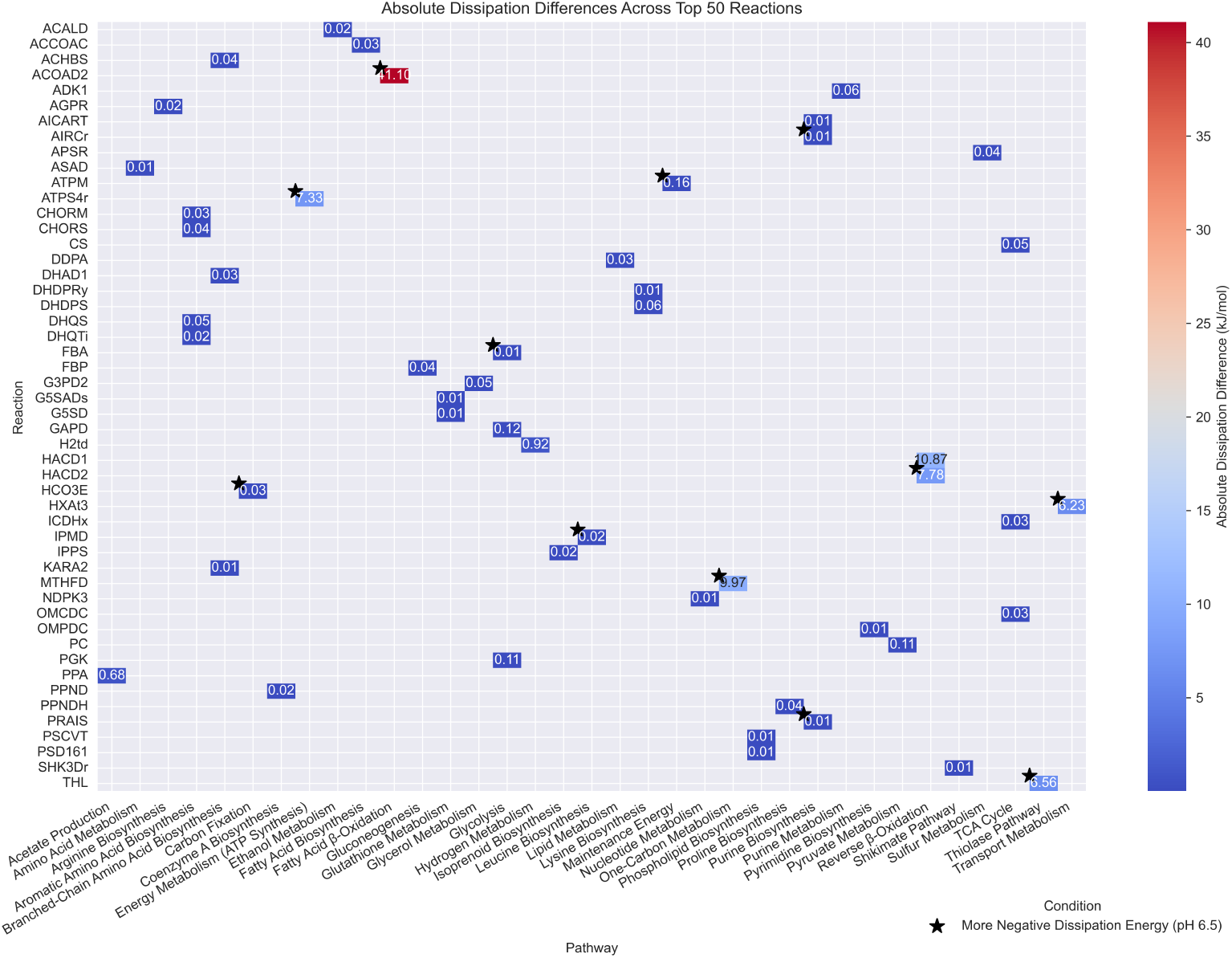
Heatmap of absolute dissipation differences across the top 50 reactions between pH 5.5 and pH 6.5. The color gradient indicates the magnitude of dissipation changes, with blue representing lower dissipation differences and red indicating higher values. The x-axis represents the pathways to which the reactions belong, while the y-axis lists individual reactions. Reactions where pH 6.5 exhibits more negative dissipation energy are marked with a black asterisk, indicating favorability under neutral conditions. The heatmap highlights both the magnitude of dissipation differences and the influence of pH on reaction favorability.

Among the top 50 reactions, HACD1 (3-hydroxyacyl-CoA dehydrogenase (acetoacetyl-CoA)) and HACD2 (3-hydroxyacyl-CoA dehydrogenase (3-oxohexanoyl-CoA)) displayed notable dissipation differences. HACD1 exhibited an 11 kJ/mol increase in dissipation at pH 6.5 compared to pH 5.5 but had a more negative dissipation energy at pH 5.5, indicating favorability under mildly acidic conditions. In contrast, HACD2 exhibited a 7.8 kJ/mol increase in dissipation at pH 6.5, with pH 6.5 having a more negative dissipation energy, highlighting its favorability under neutral conditions. Similarly, ACOAD2 (Acyl-CoA dehydrogenase) demonstrated the largest dissipation difference among all reactions analyzed, with a difference of 41 kJ/mol, emphasizing the energetic adaptation of fatty acid *β*-oxidation processes under varying pH conditions.

Interestingly, these findings show a mixed alignment with experimental observations under autotrophic conditions, where chain elongation has been reported to be more feasible at pH 5.5 than at pH 6.5 (Mariën et al., 2024). While some reactions, such as HACD1, support the experimental results by demonstrating favorability under acidic conditions, others, like HACD2, suggest enhanced favorability at neutral pH, potentially countering the experimental trends. This nuanced interplay highlights both the strengths and limitations of thermodynamic predictions in capturing the complexity of metabolic regulation, which includes enzyme activity, substrate availability, and kinetic constraints—factors not explicitly modeled in TFA. These results underscore the importance of integrative approaches that combine thermodynamic insights with experimental data to more comprehensively understand pH-driven metabolic shifts in *C. luticellarii*.

While most reactions showed moderate dissipation differences, reactions like PC (Pyruvate Carboxylase) and PGK (Phosphoglycerate Kinase) exhibited dissipation differences of approximately 0.11 kJ/mol. These differences suggest subtle energy reallocation within central carbon metabolism, fine-tuning energy use while maintaining overall stability. The findings here illustrate that reactions like HACD1, HACD2, and ACOAD2 act as flexible nodes in the metabolic network, fine-tuning their energy requirements in response to pH changes. These findings emphasize the dual role of *C. luticellarii*’s metabolism: maintaining overall stability while selectively optimizing key reactions to enhance metabolic efficiency under varying environmental conditions.

#### 2.4.1. Pathway-specific dissipation patterns

A focused view of flux distributions and energy dissipation changes across key metabolic pathways, notably the Wood-Ljungdahl Pathway (WLP) and the branched tricarboxylic acid (TCA) cycle, under pH 5.5 and 6.5 is presented (see Figure 8). The top 50 metabolic reactions analyzed include reactions from these pathways as well as others, selected based on their absolute dissipation differences between the two pH conditions, highlighting reactions with significant energetic adaptations. The WLP, integral for autotrophic growth through carbon fixation, shows minimal dissipation differences (as well as overall dissipation) in foundational reactions such as CODH_ACS (Carbon monoxide dehydrogenase/Acetyl-CoA synthase) and MTHFR5 (0.00 kJ/mol), suggesting that these carbon reduction steps remain tightly regulated regardless of pH. However, reactions like ACALD (Acetaldehyde dehydrogenase), which converts acetyl-CoA to acetaldehyde, show slight dissipation increases (0.0211 kJ/mol), reflecting a degree of metabolic flexibility in processing carbon intermediates.

**Figure 8:**
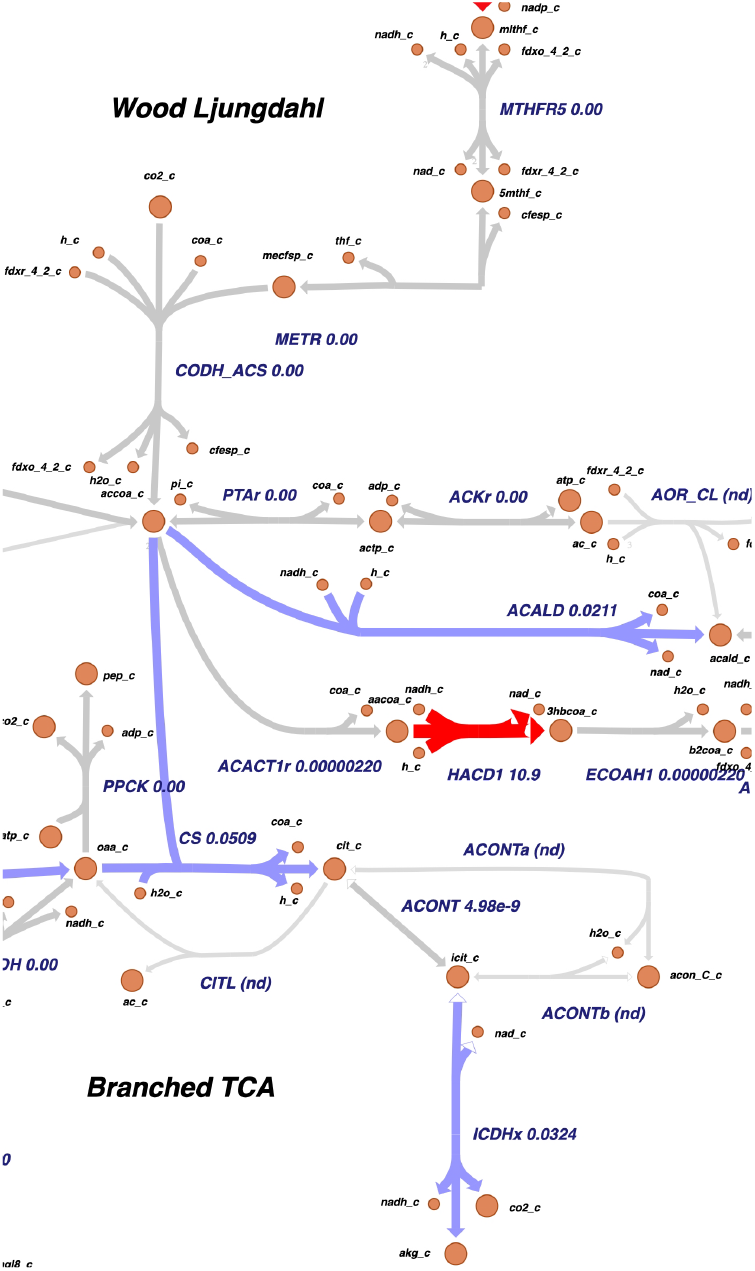
Focused view of the Wood-Ljungdahl Pathway and the branched TCA cycle with dissipation changes between pH 5.5 and pH 6.5. Red arrows indicate increased dissipation compared to pH 5.5, while blue arrows represent decreased dissipation compared to pH 5.5.

One of the more striking variations is seen in HACD1 associated reaction, where dissipation difference is 10.9 kJ/mol, indicating a significant adjustment (more negative, thus favoribility) in energy utilization when producing longer-chain compounds like acetate at a mildly acid pH of 5.5. This aligns with the role of HACD1 in energy-intensive pathways such as reverse *β*-oxidation, where the balance between pH-driven favorability and chain elongation depends on the specific metabolic context. Greater energy dissipation in a reaction often means the step becomes more thermodynamically favorable, facilitating its progression under varying conditions. However, this energy is generally dissipated, which may include heat, the diffusion of metabolites, or the release of unused energy carriers (e.g., reduced cofactors), and is not directly recoverable by the organism unless it is coupled to energy conservation mechanisms, such as electron bifurcation (e.g., co-reduction of ferredoxin) or the generation of a proton motive force, as is the case for this reaction. Without such coupling, the dissipated energy cannot be utilized for processes like H^+^ translocation or ATP synthesis. The organism instead compensates through adjustments in other metabolic pathways, highlighting the balance between flexibility and efficiency in maintaining homeostasis.

In contrast, the branched TCA cycle demonstrates a more nuanced pattern of energy adjustment. Reactions like CS (Citrate synthase), responsible for the synthesis of citrate from acetyl-CoA and oxaloacetate, show modest dissipation (0.0509 kJ/mol), suggesting some flexibility in the pathway’s entry points under varying pH (8). ICDHx (Isocitrate dehydrogenase) exhibits a dissipation of 0.0324 kJ/mol, indicating adjustments in the conversion of isocitrate to *α*-ketoglutarate, a critical step for generating reducing equivalents (NADH). These smaller dissipation changes reflect a balanced response to pH shifts, focusing on maintaining energy efficiency within the core metabolism. The observed variability in WLP and specific TCA reactions underscores how *C. luticellarii* reallocates energy based on environmental conditions, using pathways like the WLP for broader energy adjustments at pH 6.5 while keeping essential metabolic processes stable at lower pH levels. This detailed energy management strategy allows the organism to optimize its metabolic outputs, whether in carbon fixation or energy conservation.

#### 2.4.2. Variability in metabolic pathway flexibility

Thermodynamic Variability Analysis (TVA) revealed how *Clostridium luticellarii* adjusts its metabolic network across different pH conditions, shedding light on its capacity for metabolic flexibility. At pH 6.5 (see Supplementary Figure S3), the network exhibited greater flux variability, particularly in key pathways like the Wood-Ljungdahl Pathway (WLP) and reverse *β*-oxidation, both of which are crucial for autotrophic growth and the synthesis of longer-chain carboxylates such as butyrate and isobutyrate. The WLP’s increased variability in carbon fixation, specifically in reactions reducing CO_2_ to acetyl-CoA, indicates greater adaptability at pH 6.5, which directly supports higher yields of these energy-demanding products.

In contrast, TVA at pH 5.5 (see Supplementary Figure S4) showed reduced variability, especially in pathways with high ATP demand, such as WLP and the ATP-investment steps of glycolysis. The heightened proton concentration at this pH imposes greater energetic costs on ATP production, resulting in a more constrained metabolic state. This reduced flexibility limits the organism’s ability to adjust its metabolic fluxes, making it less responsive to environmental changes compared to the adaptability observed at pH 6.5.

The differences in metabolic flexibility between pH conditions significantly impact the production capabilities of *C. luticellarii*. At pH 5.5, the constrained flexibility in pathways like carbon fixation and chain elongation limits the production of higher-value products such as butyrate and isobutyrate, favoring acetate production due to its lower energy demands. In contrast, pH 6.5 allows greater metabolic flexibility, enabling a broader range of energy-intensive products by optimizing pathways like the WLP and reverse *β*-oxidation. This adaptability at pH 6.5 is advantageous for producing higher-value fermentation products, making it a preferred condition for maximizing output. The combined insights from TFA and TVA underscore how pH influences the balance between metabolic rigidity and flexibility, guiding the optimization of fermentation conditions for desired outputs.

#### 2.4.3. Integration of dissipation and variability analysis

The integration of TFA and TVA results reveals how *C. luticellarii* adjusts its metabolic strategies across different pH conditions, highlighting both its flexibility and constraints. At pH 6.5, increased energy dissipation and higher flux variability in key pathways like HACD1 indicate a more adaptable metabolic network, allowing efficient carbon fixation and chain elongation that support the production of energy-intensive products like butyrate and isobutyrate. In contrast, at pH 5.5, reduced flux variability and smaller changes in dissipation reflect a more rigid metabolic state, as the organism adapts to the energetic challenges of higher proton concentrations. This rigidity, as predicted by the model simulations, limits the production of complex carboxylates and instead favors a metabolic shift toward simpler products like acetate.

## 3. Discussion

This study presents *iSJ444*, a model for *C. luticellarii* adapted from its close relative *C. ljungdahlii*. This model enabled the evaluation of the chain elongation pathways from earlier studies and the generation of predictions for not yet experimentally tested metabolic behavior. The model is highly rated by MEMOTE and shows growth on multiple substrates relevant to syngas fermentation. Simulation results support that the earlier hypothesized pathways are most likely utilized by *C. luticellarii*. The performance of *iSJ444* also highlights the robustness of *iHN637*, despite it being relatively outdated (published in 2013). Comparisons to experimental results were made without constraining the actual uptake of metabolites. Yet, the product spectra matched well, demonstrating the power of flux sampling and the utility of metabolic models for predictions, even without precise data on substrate use. This study supports the further testing and inclusion of *C. luticellarii* in syngas-fermenting systems.

The *iHN637* model of *C. ljungdahlii* was chosen as a template for model creation, but *C. kluyveri* could have also served this purpose. A *C. kluyveri* metabolic model exists (Zou et al., 2018), and it is the next closest strain to *C. luticellarii*, with eggNOG mapping indicating many *C. luticellarii* genes are related to *C. kluyveri* (Table 1). However, its model was less accessible and conventional than the BiGG-formatted *iHN637*. The *iHN637* model also includes ferredoxin-linked reactions, such as those involving electron bifurcation and ferredoxin-NAD(P) reductase (RnF) complexes, which are key to energy conservation and are most likely present in *C. luticellarii* as well. These features made *iHN637* a practical and effective starting point for creating the *iSJ444* model. While a clostridial metamodel that consolidates key features of closely related species and can be pruned or tuned for specific organisms would be highly useful, it would require careful design to balance generality and accuracy. Such a framework could streamline future efforts to model less-characterized *Clostridia* by providing a robust starting point with established core pathways. Future improvements to *iSJ444*, such as incorporating additional reactions from *C. kluyveri*, could contribute toward developing a more universal Clostridial model, enabling broader comparisons and applications.

*iSJ444* is less extensive than similar models of related acetogens. The model has 735 reactions, whereas metabolic models of *C. kluyveri, C. autoethanogenum*, and *Clostridium tyrobutyricum* have 994, 1109, and 858 reactions, respectively (Feng et al., 2022; Valgepea et al., 2017; Zou et al., 2018). This smaller size is due to a combination of factors: i) the *iHN637* model, which served as the starting point, is itself smaller compared to some more recent models; and ii) additional reactions were removed during the adaptation process to focus on pathways relevant to syngas metabolism and chain elongation in *C. luticellarii*. By taking this reductionist approach, *iSJ444* achieves greater specificity and computational efficiency, enabling more accurate predictions of syngas fermenting and chain elongation capabilities. This focused design minimizes the inclusion of extraneous reactions that are not experimentally supported for *C. luticellarii*, reducing uncertainty and simplifying the interpretation of simulation results. Results of growth on glucose and glycerol demonstrate some of the limitations. This type of constraint-based modeling has inherent limitations; gene expression is usually not considered, though methods exist for this (Åkesson et al., 2004). The lack of experimental data on *C. luticellarii*, in particular, poses challenges, as the formulation of organism-specific biomass functions often requires extensive experimental data, which is crucial for model accuracy (Dikicioglu et al., 2015; Lachance et al., 2019). While *iSJ444* is not intended for broader exploration of *C. luticellarii*’s metabolism, its specificity makes it a powerful tool for investigating and optimizing syngas-based bioprocesses and targeted metabolite production. This design strategy highlights the trade-off between model comprehensiveness and precision, with *iSJ444* being tailored for its intended applications.

When comparing the model simulation results, it is important to note that model simulations represent fluxes at a single time point, whereas experimental results reflect changes in metabolite concentrations over the entire incubation period. The comparisons made here thus involve production/consumption at a specific time point for the model versus total production/consumption at the end of the incubation for the experiments. As production and consumption do not remain constant over time, this discrepancy should be considered when interpreting the comparisons.

The model accurately predicted the product spectrum of methanol fermentation but overestimated the formation of chain elongation products and H_2_, while CO_2_ levels were similarly elevated. This overestimation could be linked to the model’s inability to fully capture the dynamic interplay between H_2_ and CO_2_ consumption, particularly under conditions where CO_2_ serves as the primary electron acceptor. In acetogens, significant H_2_ production is uncommon when electron acceptors such as CO_2_ are available, which may explain the deviation from experimental observations. This discrepancy may arise from differences between the model’s steady-state predictions and the cumulative production in batch incubation. Dynamic conditions during incubation, such as nutrient shifts, early growth biomass production, or limitations in gas-liquid mass transfer, commonly observed in batch systems could divert resources from product formation, thereby reducing observed yields. Imposing constraints on biomass and ATP generation could potentially resolve this issue. In the simulations, propionate use was limited. In the model, only a single propionate utilization pathway was considered (valerate production). The low uptake rates could be attributed to propionate metabolism into valerate, which yields insufficient energy. When forcing a methanol/propionate uptake ratio similar to that observed in experiments, *iSJ444* predicted significantly more valerate production than was observed experimentally. As previously hypothesized, this variation may be due to the presence of yet unidentified energy generating reactions within the pathway or oxidation, suggesting that alternative pathways could be involved in the metabolism of propionate in *C. luticellarii* (De Smit et al., 2019; Mariën et al., 2024). The base *iHN637* model of *C. ljungdahlii* could form lactate from propionate, a capability that was removed in *iSJ444* to prevent the production of valerate/propionate when simulating growth on methanol, which was not observed in experiments.

Comparisons to experimental data were made more challenging due to the high standard deviations observed during flux sampling of the *iSJ444* model. Even with a large number of samples, the results do not appear to converge, potentially due to the simplicity of the added pathways or because the energy production options are closely balanced. The metabolites not present in *iHN637* and added to *iSJ444* have only two associated reactions. The chain elongation pathways added are very linear, with no alternative pathways for metabolite entry. When flux is randomly sampled, the quantity of chain elongation products is heavily dependent on the initial flux of the reaction that initiates this linear pathway. Because the inflow and outflow flux of all metabolites must be balanced, all fluxes in such a linear pathway are interdependent. Thus, standard deviations of end products may remain high even with many samples, as they depend on a single initial flux. Including reactions that produce intermediates in the pathway could reduce the standard deviation of the final products. These reactions (if present) could be identified by examining *C. kluyveri*, a more extensively studied chain elongator. Nonetheless, considering the simplicity of the *iSJ444* model and its chain elongation pathways, the model matches the experimental data on methanol and H_2_/CO_2_ qualitatively and, to some extent, quantitatively. This supports the presence of the even-chain elongation pathway proposed by Petrognani et al. (2020).

The batch feeding strategy employed in the experimental work by Petrognani et al. (2020) plays a significant role in understanding the discrepancies between model predictions and observed data. In these experiments, substrates were supplied at the start of the batch fermentation, resulting in dynamic changes in substrate availability and metabolite concentrations over time. Such fluctuations inherently contrast with the steady-state assumptions underlying the *iSJ444* model simulations, which consider constant flux distributions. To improve the alignment between model predictions and experimental data, future studies could implement continuous feeding strategies. This approach maintains substrate concentrations at steady levels, reducing temporal variability and creating conditions that more closely mimic the steady-state assumptions of the metabolic model. Additionally, dynamic Flux Balance Analysis (dFBA) could be employed to capture the temporal changes in metabolic activity observed in batch systems, thereby enhancing the predictive capabilities of the model under dynamic conditions (Foster et al., 2021; Scott et al., 2023). By considering both feeding strategies and dynamic modeling approaches, future experimental designs can bridge the gap between *in silico* predictions and experimental observations, ultimately improving the model’s utility in guiding biotechnological applications for syngas fermentation.

Predictions of performance on untested growth media indicated that the product spectrum on syngas was primarily influenced by carbon availability, with more ECCA formation when less carbon source (in the form of either CO or CO_2_) was supplied, while maintaining the same energy potential through H_2_. Limiting carbon uptake appears to increase the need for complete oxidation of the available carbon, yielding more ATP and producing longer carboxylic acids. *Butyribacterium methylotrophicum*, which can grow on syngas to produce ethanol, acetate, lactic acid, and butyrate, exhibited similar behavior, with a higher percentage of hydrogen in the feed increasing production (Heiskanen et al., 2007). *E. limosum* showed increased acetate production when supplied with H_2_/CO_2_ compared to only CO, but no butyrate was formed in either condition, making direct comparison with the ECCA-producing model more challenging. *C. ljungdahlii* also increased acetate production over ethanol under high H_2_/CO ratios (Jack et al., 2019). More in line with our results is the increased ethanol production of *C. autoethanogenum* under high H_2_, observed experimentally (Valgepea et al., 2018) and predicted through metabolic modeling (Benito-Vaquerizo et al., 2020). This suggests that high H_2_ may enable the utilization of pathways beyond acetate formation. While *C. luticellarii* has thus far only been grown on H_2_ and CO_2_, the effects of varying syngas compositions on the products of *C. luticellarii* warrant further experimental study.

Supplying methanol along with syngas did not result in increased ECCA production, even though results with methanol as the sole feed showed higher ECCA production than growth on syngas alone. The lack of improvement in the product spectrum with methanol aligns with the simulations on methanol and CO_2_, as CO_2_ led to increased acetate production. In the presence of syngas, a similar effect occurs, where uptake of CO and CO_2_ is predicted to lead to more acetate production. *E. limosum* was found to grow and take up syngas more rapidly when simultaneously fed with methanol, but the product spectrum was not measured in that study (Kim et al., 2021). It was found that the accelerating effect of methanol only occurred during growth on H_2_ and CO_2_, and not for CO and CO_2_, due to a closer connection between the H_2_/CO_2_ and methanol enzymatic pathways. Although this effect would not show in the simulations, it is important to consider for future studies on syngas and methanol metabolism by *C. luticellarii*.

Future experimental validation of the *iSJ444* model under varying syngas compositions is critical for testing and refining its predictive capabilities. Such efforts could benefit from adopting a Design-Build-Test-Learn (DBTL) cycle, a well-established framework in metabolic engineering that enables iterative improvements in both experimental and computational workflows (Nielsen and Keasling, 2016; Gurdo et al., 2022). In the “Design” phase, the *iSJ444* model can be used to simulate metabolic responses under different H_2_/CO/CO_2_ ratios, identifying conditions likely to enhance product formation or expose metabolic bottlenecks. During the “Build” and “Test” phases, these conditions can be experimentally implemented and validated, while the “Learn” phase allows integration of the results into subsequent model iterations. This iterative approach has proven effective in improving the predictive power of genome-scale metabolic models and optimizing bioprocesses (Otero-Muras and Carbonell, 2021). By leveraging the DBTL cycle, the *iSJ444* model could serve as a dynamic tool for systematically exploring syngas fermentation, facilitating the development of resilient and efficient industrial processes. Incorporating this strategy aligns with the broader aim of bridging computational predictions with experimental insights in metabolic engineering.

It is challenging to accurately represent pH in metabolic model simulations, particularly in the context of chain elongation processes (de Leeuw et al., 2020; Ganigué Pagès et al., 2016). However, with sufficient experimental data and assumptions, pH effects can be incorporated into genome-scale models to enhance predictive power. Extensive work on Thermodynamic Flux Analysis (TFA) has provided a basis for integrating thermodynamic constraints into metabolic models, effectively improving the accuracy of metabolic network simulations (Müller and Bockmayr, 2013; Gollub et al., 2021). In addition to thermodynamic constraints like pH, incorporating proteomics data with tools such as GECKOpy and ECMpy can further refine genome-scale model predictions (Carrasco Muriel et al., 2023; Mao et al., 2024). Integrating proteomics data alongside thermodynamic factors enables finer constraint adjustments, resulting in more accurate flux predictions. However, the lack of experimental proteomics data for *C. luticellarii* currently limits opportunities to further constrain our genome-scale model using these approaches.

*iSJ444* serves as a minimal yet robust base model of *C. luticellarii*, suitable for further exploration of its syngas and chain elongation capabilities. The model could be improved to allow for more accurate predictions of odd-chain elongation by expanding odd-chained metabolic pathways. It could also benefit from a tailored biomass function and a well-formulated objective function if it is to be used for dFBA. Model predictions encourage further metabolic testing of *C. luticellarii*, especially concerning the effects of H_2_/CO or CO_2_ ratios on product formation, and the addition of methanol when growing on syngas, as these model predictions seem to contradict earlier studies. A large effect of syngas composition on product formation could be detrimental, as a key advantage of microbial-based syngas fermentation could come from its insensitivity to variation in composition (Liew et al., 2016). Beyond direct use of syngas, *C. luticellarii* might be more productive when supplied with syngas-derived methanol, and optimization of this approach can also be studied using this model. Insights from the chain elongation pathways of *C. kluyveri* could be leveraged to further enhance *iSJ444*. The model can also be integrated into consortia, including those with *C. autoethanogenum*, a popular organism in syngas fermentations that has an established metabolic model (Valgepea et al., 2017). As *C. autoethanogenum* has similar metabolic properties to *C. ljungdahlii*, it would likely play a similar role in a consortium model. This study represents an initial step in exploring the potential of *C. luticellarii* for enhancing syngas recycling and suggests further experimental and *in-silico* testing of its capabilities.

## 4. Materials and methods

### 4.1. Model creation: *iSJ444*

The genome-scale metabolic model (GEM) for *Clostridium luticellarii*, named *iSJ444*, was constructed using a combination of ortholog mapping and manual curation. The *iHN637* model of *Clostridium ljungdahlii* (Nagarajan et al., 2013), retrieved from the BiGG database (http://bigg.ucsd.edu/models/iHN637), was used as the base model. Orthologous genes were identified using the eggNOG-mapper tool (Cantalapiedra et al., 2021), and reactions corresponding to genes not present in the *C. luticellarii* genome were removed from *iHN637*. Reactions that were orphaned by this process but had single Enzyme Commission (E.C.) annotations were reinstated if orthologous genes performing the same E.C. function were identified in the *C. luticellarii* genome.

To ensure the model’s functionality, MEMOTE analysis (Lieven et al., 2020) was employed to detect and fill any pathway gaps, allowing the model to produce all essential biomass precursors. Specific chain elongation pathways, relevant for butyrate and isobutyrate production, were manually incorporated based on previously published metabolic characterizations of *C. luticellarii* (De Smit et al., 2019; Petrognani et al., 2020). Where possible, reaction identifiers from the BiGG database were used. Finally, flux bounds were adjusted to fit experimental data, ensuring that the model’s predictions aligned with the observed growth and product formation conditions. Additionally, FROG (Flux Robustness and Optimization for GEMs) analysis (Raman et al., 2024) was performed to confirm the reproducibility and robustness of the flux balance analysis (FBA) (Orth et al., 2010) modeling results.

The setup of this study, showing how the *iHN637* model of *C. ljungdahlii* was adapted to the *iSJ444* model for *C. luticellarii* using orthology gene mapping (eggNOG-mapper) and the addition of hypothesized chain elongation pathways (De Smit et al., 2019; Petrognani et al., 2020) is depicted in a diagram (see Figure 1). *iSJ444* was created with the potential to be incorporated into consortia models along with other preexisting metabolic models to assess their potential for syngas recycling.

### 4.2. Model validation conditions

The validation of the *iSJ444* GEM against experimental data was conducted using results from batch fermentation experiments as described by Petrognani et al. (2020). In these experiments, *Clostridium luticellarii* DSM 29 923 was cultivated on a synthetic methanol medium supplemented with 200 mM methanol as the electron donor, 100 mM potassium acetate, and 23 mM sodium butyrate as electron acceptors. The experiments were conducted in sealed, static penicillin bottles under controlled conditions, where substrates were added at the start of incubation rather than through continuous feeding. The ‘average electron equivalent’ for the product spectrum was calculated by multiplying the flux of each product by its degree of reduction, as listed in Supplementary Table S1. The resulting values were normalized against the total electron equivalents of all products in the spectrum. This calculation method was applied to both experimental and model-predicted data to enable direct comparisons.

To provide an accessible summary of the experimental data used for validation, a table has been provided outlining the experimental sources, conditions, and key findings (Table S3). This table highlights the limited, but critical datasets used for validating the *iSJ444* GEM and emphasizes the importance of these data in guiding model predictions and interpretations.

### 4.3. Flux sampling simulations

The genome and amino acid sequences of *C. luticellarii* (strain PVXP01) were retrieved from the NCBI sequence database (Poehlein et al., 2018). Model simulations were performed using COBRApy version 0.24.0 (Ebrahim et al., 2013) with CPLEX optimizer version 22.1.0.0 as the solver, running in Python 3.8. Unless otherwise specified, flux sampling was conducted using COBRApy’s built-in artificial centering hit-and-run (ACHR) algorithm with 15,000 samples, based on the sampling strategy used in previous syngas-fermenting co-culture studies (Benito-Vaquerizo et al., 2020). Flux sampling was chosen for its ability to generate diverse, non-optimal flux distributions, allowing for a more realistic representation of metabolic variability compared to methods that optimize a single objective (Herrmann et al., 2019). Optimized General Parallel Sampler (OPTGP) sampling was also tested, but no significant differences were observed compared to ACHR sampling. Flux variability analysis was also performed on *iSJ444* using COBRApy, with the loopless setting on true and the fraction of optimum on 0, to see the full solution space (Table S4).

Flux sampling was employed to simulate the metabolic activity of *C. luticellarii* under various conditions. Growth simulations did not rely on maximizing the biomass reaction but instead used flux sampling to explore a range of possible metabolic flux distributions. While the biomass reaction provides a theoretical framework for cellular growth, it was not used as an optimization objective in this study, aligning with the study’s focus on product spectrum predictions rather than growth yield.

### 4.4. Thermodynamic flux analysis and energy dissipation

Thermodynamic flux analysis (TFA) (Henry et al., 2007) was conducted using the pyTFA Python package (Salvy et al., 2019), which incorporates thermodynamic constraints into flux balance models to improve the realism of metabolic predictions. We adapted pyTFA tutorials on sampling (Salvy et al., 2019) and Equilibrator integration (Beber et al., 2022) to perform TFA, thermodynamic variability analysis, and flux sampling, allowing us to assess energy dissipation across the metabolic network. The use of Equilibrator (Beber et al., 2022) allowed us to incorporate accurate thermodynamic data, improving the precision of reaction feasibility predictions by ensuring that Gibbs free energy calculations were context-specific and consistent with the intracellular environment.

Due to the lack of specific thermodynamic data for *C. luticellarii*, assumptions were made regarding intracellular compartment conditions at pH 5.5 and pH 6.5. In particular, compartment pH values and the membrane potential were inferred from similar organisms, as no experimental data are available for *C. luticellarii* under these conditions. These assumptions were essential for setting boundary conditions for TFA simulations. By integrating Gibbs free energy data and simulating growth under both acidic (pH 5.5) and neutral (pH 6.5) conditions, we quantified the energy dissipation and identified potential metabolic bottlenecks that impact the efficiency of key pathways. Flux sampling was used to capture the distribution of feasible metabolic states, providing insights into the robustness of metabolic pathways related to butyrate, isobutyrate, and acetate production.

## 5. Abbreviations

### 5.1 Feedstocks

In brackets are influx rates/ratios used in the simulations/experiments.

MA: Methanol + Acetate (20/10)
MAB: Methanol + Acetate + Butyrate (20/10/2.5)
MP: Methanol + Propionate (20/10)
MPV: Methanol + Propionate + Valerate (20/10/2.5)
LA: D-Lactate + Acetate (20/10)
GLC: Glucose (2.5)
GLY: Glycerol (2.5)
H2-CO2: Hydrogen and carbon dioxide (80/20)
Syngas: Unless otherwise specified: hydrogen, carbon monoxide, and carbon dioxide in a 30/60/10 ratio
ME: Methanol (20)
MCO2: Methanol and carbon dioxide (20/10)

### 5.2 Model metabolite names

_c suffix means intracellular metabolite, _e extracellular.

meoh: Methanol
ac: Acetate
ppa: Propionate (C3)
but: Butyrate (C4)
ibt: Isobutyrate (C4)
pta: Valerate (C5)
hxa: Caproate (C6)
accoa: Acetyl-CoA
aacoa: Acetoacetyl-CoA
fdxr/o_42: Ferredoxin reduced/oxidized form 4:2

### 5.3. Model reaction names

MTAabcPS: Pseudo reaction of the methanol-cobalamin methyltransferase system
CODH4: Carbon monoxide dehydrogenase
METR: Methyltetrahydrofolate:corrinoid/iron-sulfur protein methyltransferase
CODH_ACS: Carbon monoxide/acetyl-CoA synthase pseudo reaction
VCACT: Acetyl-CoA C-acyltransferase
HVCD: 3-hydroxyacyl-CoA dehydrogenase
VECOAH, ECOAH1 and ECOAH2: 3-hydroxyacyl-CoA dehydratase
VCOAD2, ACOAD1z, ACOAD2: Acyl-CoA dehydrogenase
FACOAL50i: Fatty acid CoA ligase
ACACT1r: Acetyl-CoA C-acetyltransferase
HACD1, HACD2: 3-hydroxyacyl-CoA dehydrogenase
BUTCT: Acetyl-CoA:butyrate-CoA transferase
IBTMr: Isobutyryl-CoA mutase
IBUTCT: Acetyl-CoA:isobutyrate-CoA transferase
THL: Thiolase
HXCT: Acetyl-CoA:hexanoate-CoA transferase

## 6. Acknowledgments

W.T.S.J, P.J.S, and J.J.K acknowledge the Dutch Research Council (NWO), and Wageningen University & Research for their financial contribution to the Unlock initiative (NWO: 184.035.007).

## 7. Data & code availability

**Genome-scale models**: The GEM as well as accompanying MEMOTE (Lieven et al., 2020) and FROG Analysis (Raman et al., 2024) reports can be found in a public repository on BioModels: https://www.ebi.ac.uk/biomodels/. Fo the *iSJ444* GEM, see here: MODEL2411260001. **Simulation code**: The code used for the model simulations can be found at https://gitlab.com/wurssb/Modelling/Clostridium_luticellarii **Project data**: The study’s *in silico* data can be found on Zenodo: 10.5281/zenodo.14221978

## CRediT authorship contribution statement

**William T. Scott:** Visualization, Validation, Methodology, Investigation, Formal analysis, Supervision, Conceptualization, Writing – original draft, Writing - Proofreading and editing. **Siemen Rockx:** Visualization, Validation, Methodology, Investigation, Formal analysis, Conceptualization, Writing – original draft. **Quinten Mariën:** Data curation, Conceptualization, Writing - Proofreading and editing. **Alberte Regueira:** Data curation, Writing - Proofreading and editing. **Pieter Candry:** Conceptualization, Writing - Proofreading and editing. **Ramon Ganigué:** Data curation, Project administration, Writing - Proofreading and editing. **Jasper J. Koehorst:** Supervision, Writing - Proofreading and editing. **Peter J. Schaap:** Project administration, Funding acquisition, Writing - Review and editing.

