## Supplementary material for "Implementation of a *Clostridium luticellarii* genome-scale model for upgrading syngas fermentations": Table S3

**Table S3.** Overview of experimental data used for validating the *iSJ444* model. Each dataset includes information on the experimental conditions and the key findings utilized in the model validation.

| Study | Reactor Setup | Culture Conditions | Substrate Conditions | Products Measured | Alignment with model | Reference |
|---|---|---|---|---|---|---|
| **Petrognani et al. (2020)** | Batch | Monoculture | Methanol + Acetate | Butyrate, Isobutyrate, Caproate | Modeled spectrum aligned with experimental data | Petrognani et al., 2020 |
| **Mariën et al. (2024)** | Batch | Monoculture | H2 + $CO_2$ | Acetate, Butyrate | Deviations observed in propionate utilization | Mariën et al., 2024 |
| **De Smit et al. (2019)** | Batch & Continuous anaerobic bioreactor | Mixed culture | Methanol + Propionate | Valerate | Over-prediction of valerate in model | De Smit et al., 2019 |
