## Supplementary material for "Implementation of a *Clostridium luticellarii* genome-scale model for upgrading syngas fermentations": Table S4

*Table S3. Flux variability analysis and mean and standard deviation of flux sampling of iSJ444. MA is methanol and acetate (20 and 10 mmol/gDdW/h), MP: methanol and propionate (20 and 10 mmol/gDW/h), and H2-CO2 (80 and 20 mmol/gDW/h). Units are mmol/gDW/h. Minimum and maximum values were calculated using the built in function flux_variability_analysis of COBRApy, with the loopless setting on true and the fraction of optimum on 0, to see the full solution space.*

| | | METHANOL | ACETATE | PROPIONATE | ISOBUTYRATE | BUTYRATE | VALERATE | CAPROATE | HYDROGEN | $CO_2$ |
|---|---|---|---|---|---|---|---|---|---|---|
| **MA** | Min | -20 | -10 | 0 | 0 | 0 | 0 | 0 | 0 | 0 |
| | Max | -0.9 | 10 | 0 | 10 | 10 | 0 | 6 | 65 | 22 |
| | Mean (SD) | -4.92 (0.08) | -1.35 (1.08) | 0.0 (0.0) | 0.75 (0.53) | 0.73 (0.52) | 0.0 (0.0) | 0.12 (0.12) | 1.58 (1.09) | 0.12 (0.11) |
| **MP** | Min | -20 | 0 | -10 | 0 | 0 | 0 | 0 | 0 | 0 |
| | Max | -0.9 | 10 | 0 | 5 | 5 | 10 | 3 | 60 | 20 |
| | Mean (SD) | -4.92 (0.08) | 0.55 (0.39) | -0.14 (0.14) | 0.26 (0.19) | 0.27 (0.2) | 0.14 (0.14) | 0.09 (0.08) | 3.31 (0.52) | 0.11 (0.1) |
| **H2-CO2** | Min | 0 | 0 | 0 | 0 | 0 | 0 | 0 | -55 | -20 |
| | Max | 0 | 10 | 0 | 5 | 5 | 0 | 3 | -2 | -1 |
| | Mean (SD) | 0.0 (0.0) | 2.55 (1.83) | 0.0 (0.0) | 1.22 (0.85) | 1.14 (0.83) | 0.0 (0.0) | 0.36 (0.33) | -45.88 (2.1) | -19.65 (0.34) |
