## Supplementary material for "Implementation of a *Clostridium luticellarii* genome-scale model for upgrading syngas fermentations": Table S1

*Table S1. Degrees of reduction of products of interest, used for calculating electron equivalent product spectra in figures.*

| **Metabolite** | **Name in model** | **Degree of reduction** |
|---|---|---|
| Methanol | meoh_e | 6 |
| Ethanol | etoh_e | 12 |
| Acetate | ac_e | 8 |
| Propionate | ppa_e | 14 |
| Isobutyrate | ibut_e | 20 |
| Butyrate | but_e | 20 |
| Isovalerate | ival_e (not present in models) | 26 |
| Valerate | pta_e | 26 |
| Caproate | hxa_e | 32 |
| Hydrogen | h2_e | 2 |
| Carbon monoxide | co_e | 2 |
