## Supplementary material for "Implementation of a *Clostridium luticellarii* genome-scale model for upgrading syngas fermentations": Table S2

*Table S2. Changes in reaction bounds from iHN637 to iSJ444. Bounds are described as a minimal and maximal flux value allowed through the reactions.*

| NAME | REACTION | BOUNDS IHN637 | BOUNDS ISJ444 | REASON FOR CHANGE |
|---|---|---|---|---|
| **FTHFLI** | atp_c + for_c + thf_c --> 10fthf_c + adp_c + pi_c | 0, 1000 | -1000, 1000 | Reversible to allow ATP generation from methanol |
| **ALCD2X** | etoh_c + nad_c <=> acald_c + h_c + nadh_c | -1000, 1000 | 0, 0 | KO ethanol pathway |
| **ETOHT** | etoh_e <=> etoh_c | -1000 | 1000 | KO ethanol pathway |
| **ACACT1R** | 2.0 accoa_c --> aacoa_c + coa_c | 0, 0 | -1000, 1000 | Part of chain elongation |
| **ACOAD1Z** | b2coa_c + fdxo_42_c + 2.0 nadh_c --> btcoa_c + fdxr_42_c + 2.0 nad_c | 0,0 | 0, 1000 | Part of chain elongation |
| **BUTT** | but_e --> but_c | 0,0 | -1000, 1000 | Part of chain elongation |
| **ECOAH1** | 3hbcoa_c --> b2coa_c + h2o_c | 0, 0 | -1000, 1000 | Part of chain elongation |
| **HACD1** | aacoa_c + h_c + nadh_c --> 3hbcoa_c + nad_c | 0 | -1000, 1000 | Part of chain elongation |
| **OBTFL** | 2obut_c + coa_c --> for_c + ppcoa_c | -1000, 1000 | 0,0 | Prevents lactate formation |
| **LDH_D** | | -1000, 1000 | 0, 1000 | Prevents lactate formation |
