## Supplementary material for "Implementation of a *Clostridium luticellarii* genome-scale model for upgrading syngas fermentations": Figure S1

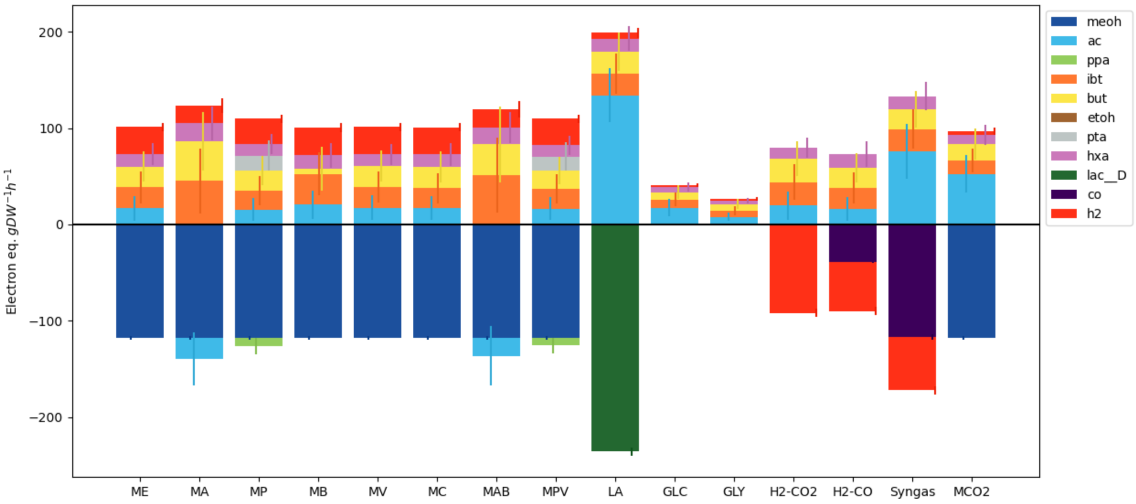


*Figure S1. Growth spectrum of C. luticellarii as predicted by iSJ444. Media are from* (Petrognani et al., 2020), *with the addition of Syngas, H2-$CO_2$, and methanol and $CO_2$. Deviation bars indicate standard deviation.*
