## Supplementary material for "Implementation of a *Clostridium luticellarii* genome-scale model for upgrading syngas fermentations": Figure S2

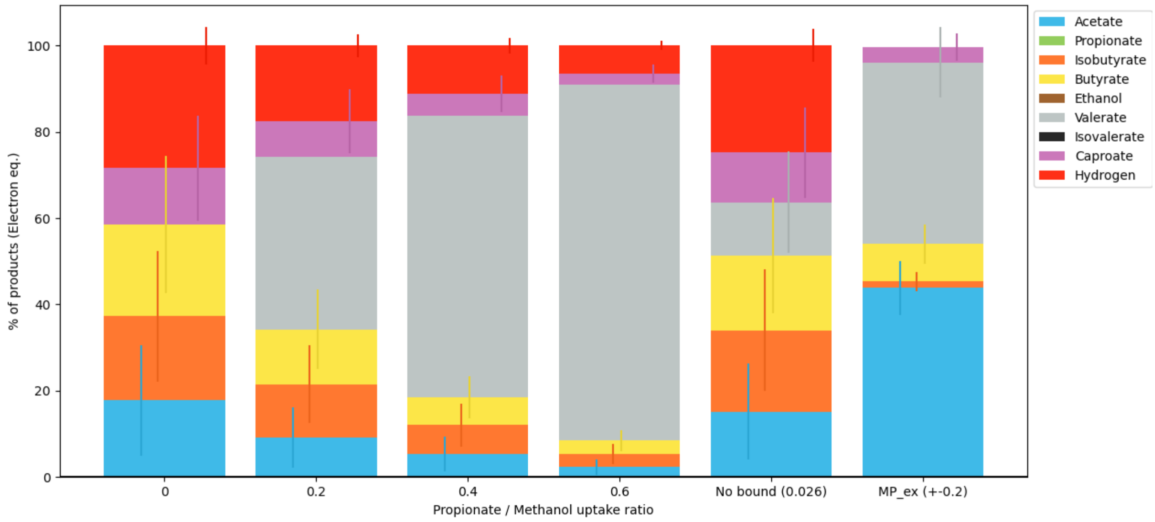


*Figure S2. Product spectrum of C. luticellarii as predicted by iSJ444 on methanol and propionate. Deviation bars indicate standard deviation.*
