## Supplementary material for "Implementation of a *Clostridium luticellarii* genome-scale model for upgrading syngas fermentations": Figure S4

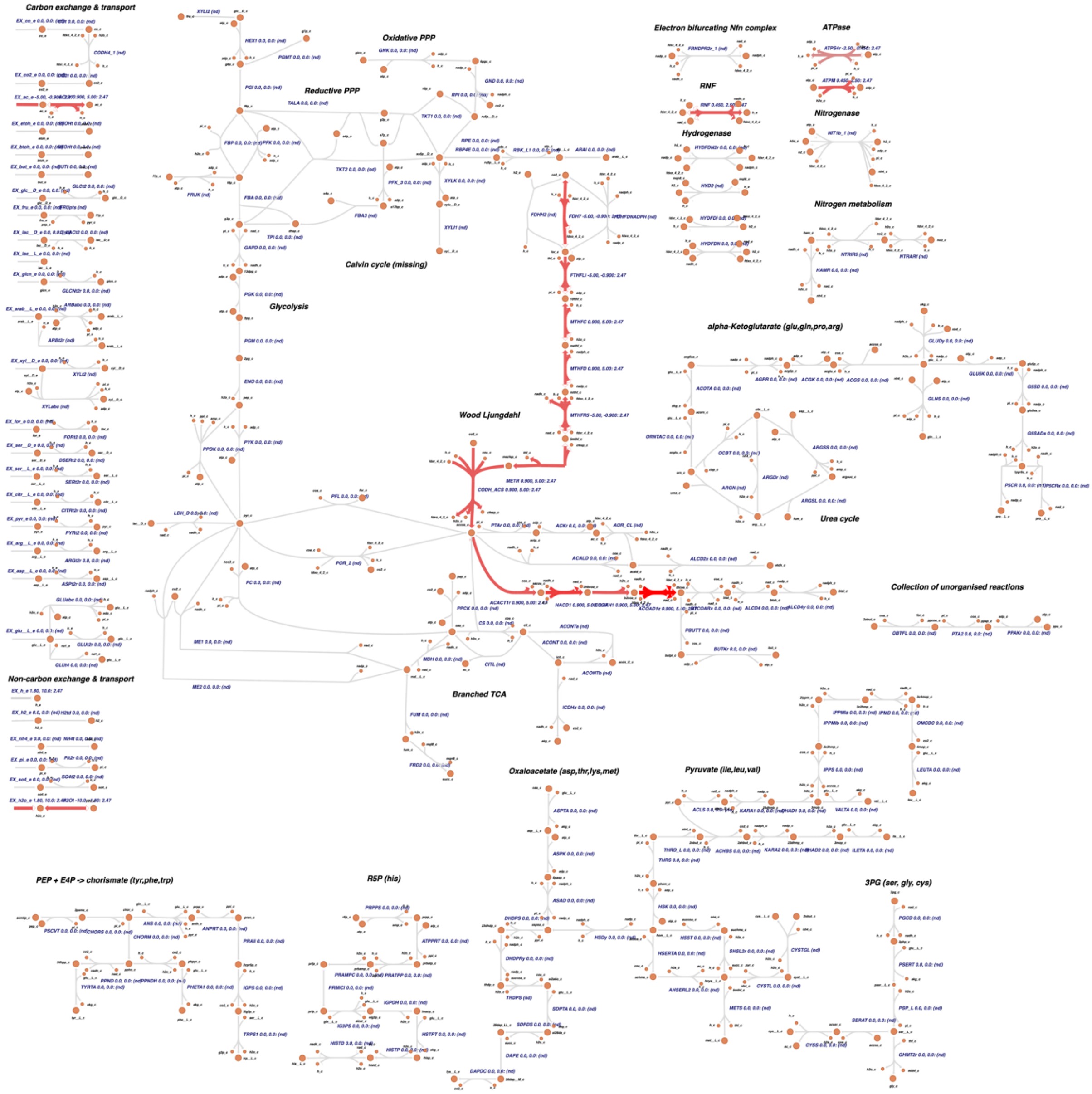


*Figure S4. Flux map showing the Thermodynamic Variability Analysis results as predicted by iSJ444 under pH 5.5. Values are in kJ/mol.*
